# Functionally redundant formate dehydrogenases enable formate-dependent growth in *Methanococcus maripaludis*

**DOI:** 10.1101/2023.05.09.540023

**Authors:** Mohd Farid Abdul Halim, Dallas R Fonseca, Thomas D Niehaus, Kyle C Costa

## Abstract

Methanogens are essential for the complete remineralization of organic matter in anoxic environments. Most cultured methanogens are hydrogenotrophic, using H_2_ as an electron donor to reduce CO_2_ to CH_4_, but in the absence of H_2_ many can also use formate. Formate dehydrogenase (Fdh) is essential for formate oxidation, where it transfers electrons for reduction of coenzyme F_420_ or to a flavin-based electron bifurcating reaction catalyzed by heterodisulfide reductase (Hdr), the terminal reaction of methanogenesis. How these competing reactions are coordinated is unknown. Furthermore, methanogens that use formate encode at least two isoforms of Fdh in their genomes, but how these different isoforms participate in methanogenesis is also unknown. Using *Methanococcus maripaludis*, we undertook a biochemical characterization of both Fdh isoforms involved in methanogenesis. Both Fdh1 and Fdh2 interacted with Hdr to catalyze the flavin-based electron bifurcating reaction, and both reduced F_420_ at similar rates. F_420_ reduction preceded flavin-based electron bifurcation activity for both enzymes. In a Δ*fdh1* mutant background, a suppressor mutation was required for Fdh2 activity. Genome sequencing revealed that this mutation resulted in loss of a specific molybdopterin transferase (*moeA*), allowing for Fdh2-dependent growth. This suggests that both isoforms of Fdh are functionally redundant, but their activities *in vivo* may be limited by gene regulation under different growth conditions. Together these results expand our understanding of formate oxidation and the role of Fdh in methanogenesis.

## Introduction

Methanogenic archaea (methanogens) are the primary producers of methane gas on Earth and are ubiquitous in anoxic environments (1–5). While methane is a potent greenhouse gas that contributes to global warming (6), it is also an attractive fuel for enhancing the efficiency of wastewater treatment processes (7–9). Most isolated methanogens are hydrogenotrophs, capable of oxidizing H_2_ as an electron donor for methanogenesis, but many can also use formate as an alternate electron donor. Compared to pathways of H_2_ oxidation, formate metabolism is poorly studied.

Formate is an end product of fermentation by many bacteria in anoxic environments where methanogens are found, and the ability of methanogens to use it as a substrate would provide a competitive advantage (10–16). Recent studies suggest that formate plays an underappreciated role as an electron donor for methanogenesis in natural environments. For example, in several members of the *Methanomicrobiales*, formate is the sole electron donor for heterodisulfide reductase (Hdr), the enzyme catalyzing the final step of methanogenesis (17, 18). In *Methanococcus maripaludis*, formate serves as an alternate electron donor when H_2_ is unavailable (19) and both hydrogenases and formate dehydrogenase (Fdh) interact with Hdr as the electron donating enzyme for the flavin-based electron bifurcating reduction of ferredoxin (for the CO_2_-reducing step of methanogenesis) and the heterodisulfide CoM-S-S-CoB (Figure 1) (20, 21). Fdh must also transfer electrons to coenzyme F_420_ (22), which is essential for the two intermediate reductions of methanogenesis. Furthermore, Fdh catalyzed F_420_ reduction is important for cells to produce H_2_ via a formate-hydrogen lyase activity that is essential for biosynthetic reaction (23–25). Fdh from *Methanobacterium formicium* contains a molybdopterin cofactor (25); however, in certain other organisms, Fdh may incorporate tungsten instead of molybdenum (26–29).

**Figure 1:**
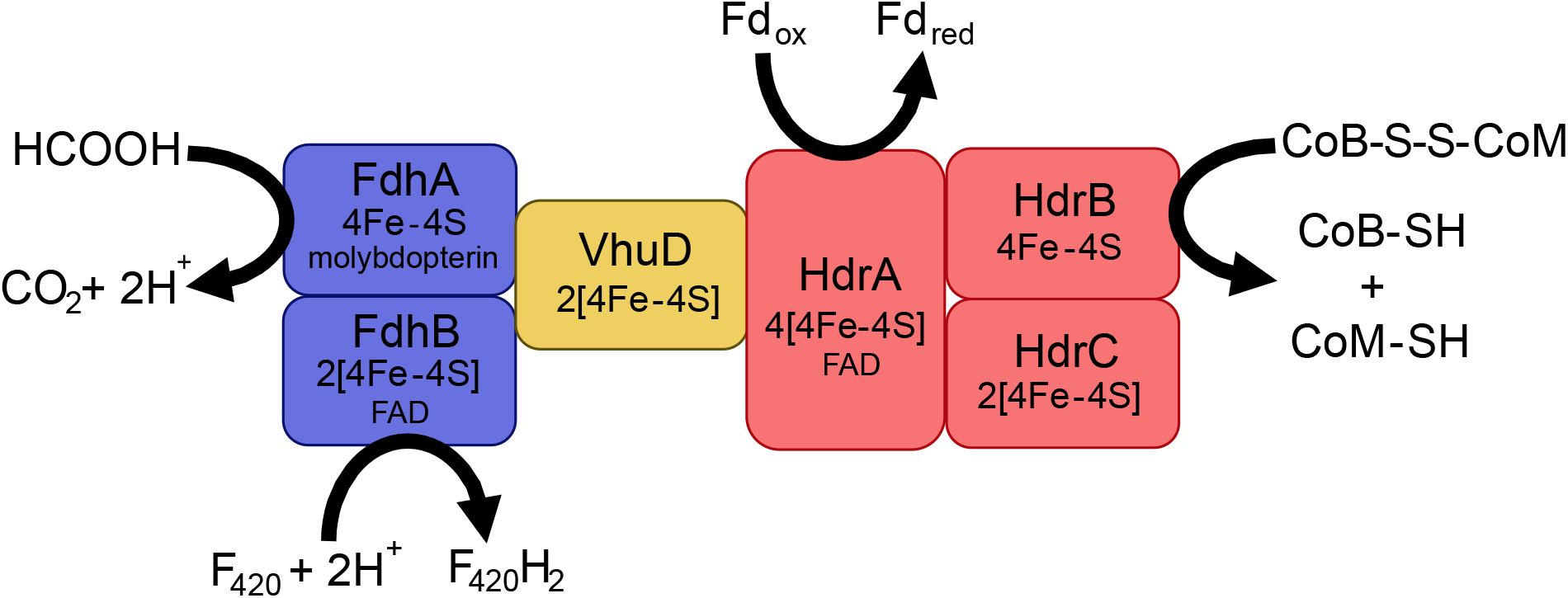
Formate dehydrogenase oxidizes formate, reduces F_420_, and donates electrons to Hdr for ferredoxin reduction via electron bifurcation.

To our knowledge, all methanogens that grow with formate encode multiple isoforms of Fdh, but whether these enzymes are responsible for different cellular functions is unclear. *M. maripaludis* encodes two Fdh homologs, *fdh1* and *fdh2*, on its genome. These were suggested to have originated from a gene duplication event (19, 30). In batch cultures, Fdh1 is critical for formate-dependent growth while Fdh2 is dispensable. Interestingly, after prolonged incubation of a τ1*fdh1* mutant strain, growth, formate oxidation, and methanogenesis occur, suggesting that a suppressor mutation allows for Fdh2-dependent growth (19, 23, 31). However, the nature of this suppressor mutation was not investigated.

To determine the roles that different isoforms of Fdh play in methanogenesis, we focus on characterizing formate oxidation in *M. maripaludis*. *M. maripaludis* is an ideal model for studying the functional redundancy of Fdh isoforms because it is genetically tractable and has a relatively simple formate metabolism with only two copies of Fdh encoding genes (19, 32). We found that both Fdh1 and Fdh2 catalyze the same reactions with comparable activity. We also characterized Δ*fdh1* suppressor mutants and found that specific molybdopterin transferases (*moeA*) appear to interact with each isoform. Combined with previous data on the regulation of *fdh* (31), this suggests that multiple isoforms of Fdh are functionally redundant with regard to activity but may be optimized for expression and activity under certain growth conditions. A deeper understanding of the mechanistic basis of formate metabolism will expand our understanding of the activity of methanogenic archaea in anoxic environments.

## Materials and Methods

### Strains, medium, and growth conditions

Strains used in this study are listed in Table S1. *M. maripaludis* S2 and its mutants were grown anaerobically at 37°C with liquid or solid McCas or liquid McCas-formate medium as previously described (33, 34). For liquid McCas medium, the following components are added (final concentration): 5 g l^-1^ NaHCO_3_, 22 g l^-1^ NaCl, 0.5 g l^-1^ NH_4_Cl, 0.34 g l^-1^ KCl, 2.8 g l^-1^ MgCl_2_·6H_2_O, 3.5 g l^-1^ MgSO_4_·6H_2_O, 0.14 g l^-1^ CaCl_2_·2H_2_O, 0.07 g l^-1^ K_2_HPO_4_, 0.0095 g l^-1^ FeSO_4_, 4.8 g l^-1^ NaCH_3_COO, 2 g l^-1^ casamino acids, and 0.001 g l^-1^ resazurin. For liquid McCas-formate medium, the following modifications were made to the McCas medium components, final concentration: 10.5 g l^−1^ NaCl, 41.8 g l^−1^ 3-(N-morpholino)propanesulfonic acid (MOPS) (pH 7), and 13.6 g l^−1^ sodium formate. To remove oxygen and any dissolved gases, the liquid medium was boiled under a stream of N_2_:CO_2_ (80:20) followed by addition of 0.5 g dithiothreitol as reductant. The medium was brought into a Coy anaerobic chamber (headspace with 2% to 3% H_2_, 10% CO_2_, balance N_2_), where 5 ml was aliquoted into Balch tubes. The headspace of Balch tubes containing liquid McCas was vacuum exchanged to H_2_:CO_2_ (80:20) and those containing McCas-formate were vacuum exchanged to N_2_:CO_2_ (80:20) to 103 kPa before autoclaving. For solid McCas/McCas-formate medium, a final concentration of 15 g l^-1^ noble agar and 2 g l^−1^ NaHCO_3_ was used. After autoclaving, the medium was cooled under a stream of N_2_:CO_2_, brought into the Coy anaerobic chamber, and poured into petri dishes.

For growth in liquid McCas or McCas-formate, 0.1 ml of a 0.5% (NH_4_)_2_S solution was added to all tubes immediately prior to inoculation. For growth on solid medium, 15 ml of 25% Na_2_S was added to an empty petri dish containing paper towels and included with the petri dishes inside an anaerobic incubation vessel. Prior to incubation, the Balch tube containing the inoculum was pressurized with their corresponding gas mix at 280 kPa. For the solid medium, the anaerobic incubation vessel was pressurized with the corresponding gas mix at 138 kPa.

### Plasmid construction

The plasmids and oligonucleotides used to construct the recombinant plasmids are listed in Table S2. All PCR fragments were generated using Phusion High-Fidelity DNA polymerase (New England Biolabs). The recombinant plasmids were assembled by T4 ligase or Gibson assembly with enzymes/reagents from New England Biolabs (35). These assembled recombinant plasmids were then transformed into *Escherichia coli* and colonies carrying the recombinant plasmid were selected as previously described (33). The recombinant plasmid constructs were verified by whole plasmid sequencing service at Plasmidsaurus.

For generating the plasmid expressing *M. maripaludis fdh2* with a C-terminal 6x-histidine tag, two PCR fragments were initially generated. Briefly, the *fdhA2* gene was PCR amplified from the start codon to the last codon encoding amino acid using the reverse primer that introduced the 6x-histidine tag. We also included the SECIS-containing DNA fragment, which encompass about 600 bp upstream of *fdh2* start codon to ensure proper translation of selenocysteine of the *fdh2* gene. Hence, a 2654 bp DNA fragment containing the *fdh2* SECIS element and *fdhA2* gene was PCR amplified. Separately, the *fdhB2* DNA fragment was also PCR amplified. The SECIS-containing DNA fragment including both *fdhA2* and *fdhB2* was assembled into *NsiI* and *AscI*-digested pLW40neo via Gibson assembly and transformed to *E. coli* DH5α via electroporation. Colonies grown on lysogeny broth solidified with 1.5% agar and ampicillin were screened by PCR to identify those carrying the correct recombinant plasmid construct.

To generate the plasmid expressing *fdh1*, we attempted a similar approach to that of the *fdh2* recombinant construct. However, the construct of *fdh1* containing a 300 bp DNA fragment upstream its gene, which contains the SECIS element, did not restore the growth of the Δ*fdh1* Δ*fdh2* strain after overnight incubation in McCas-formate. We postulated that, unlike the *fdh2* construct, the inclusion of the SECIS element for *fdh1* is not sufficient for proper expression/translation of the *fdh1* gene. Next, we generated recombinant pLW40neo carrying the *fdh1* gene without its SECIS element and compensated this by substituting the codon encoding selenocysteine to cysteine. Briefly, two fragments were generated by PCR. The first fragment was 396 bp in length, covering the *fdhA1* start codon to the new cysteine codon that was introduced by the reverse primer. The second fragment of 2831 bp was composed of the rest of the *fdhA1*. These fragments were added to *NsiI*- and *AscI-*digested pLW40neo via Gibson assembly.

The recombinant plasmid was used as a template to introduce a histidine tag at the C-terminus of *fdhA1*. Briefly, two PCR products were generated. *fdhA1* containing the cysteine substitution was amplified with a reverse primer containing the histidine tag and *fdhB1* was amplified with primers to recover the wildtype sequence. These two fragments were Gibson-assembled into *NsiI-* and *AscI*-digested pLW40neo followed by transformation and screening as described above.

### Transformation of *M. maripaludis* S2

The plasmids used in this study were transformed into *M. maripaludis* S2 Δ*upt* under anoxic conditions using a polyethylene glycol (PEG)-mediated transformation protocol (36). Briefly, *M. maripaludis* cultures were grown to late log phase with an OD_600_ of ∼0.9. The cell pellet was harvested by centrifugation and washed with transformation buffer (TB) (50 mM Tris, 350 mM sucrose, 380 mM NaCl, 1 mM MgCl_2_, pH 7.5). The washed cell pellet was resuspended in 0.375 ml of TB followed by addition of ∼5 µg of plasmid DNA. Next, 0.225 ml of PEG solution (40% wt/vol PEG 8000 in TB) was added to the cell mix followed by a 1-hour non-shaking incubation at 37 °C. Transformants were washed with fresh McCas medium and pressurized to 280 kPa with an H_2_:CO_2_ (80%:20%) gas mix. After outgrowth incubation at 37°C with agitation for at least 4 hours, transformants were sub-inoculated and selected by growth in McCas medium containing neomycin (1 mg ml^-1^)

### F_420_ purification

Ten grams (dry weight) of *M. maripaludis* cell pellet was removed from the -80°C freezer and resuspended in 15 ml of 50 mM potassium phosphate buffer (pH7). The suspended cells were boiled for 10 minutes, and the cellular debris removed by centrifugation at 14,000 × g for 20 minutes. The cleared lysate was mixed with the pre-swollen QAE-Sephadex A-25 resin equilibrated with 0.2 M NaCl in 50 mM sodium phosphate (pH 7) and allowed to incubate overnight at 4°C. The cell lysate-resin mixture was added to a disposable plastic column (1×5 cm) (Thermo Scientific) and allowed to settle. The flow-through was removed and F_420_ was eluted with a step gradient of 0 M, 0.2 M, 0.4 M, 0.6 M, 0.8 M, and 1 M NaCl (4 ml each) in the same buffer with gravity flow. The elution phase was collected in 2 ml fractions. The fraction(s) containing the eluted F_420_ were determined by measuring their absorbance at 420 nm using a Spectramax M2e microplate reader (Molecular Devices). The fractions containing the highest absorbance were pooled and pH adjusted to pH 4.7 with 1M HCl. Using a LC-10AT HPLC instrument (Shimadzu), the pooled sample was loaded onto a ZORBAX Eclipse XDB C18 column (5 μm, 4.6 × 150 mm) (Agilent) that had been equilibrated with 25 mM sodium acetate, pH 4.7. F_420_ was eluted with a 0–80% methanol gradient at a flow rate of 1 ml min^-1^ for 20 minutes. One ml fractions were collected based on the absorbance peak at 420 nm. The collected fractions were pooled and adjusted to pH 7 with 1M NaOH and run on a DEAE Sepharose column (1×5 cm). The concentration of F_420_ was calculated with the following extinction coefficient (ε420=38.5 mM^-1^ cm^-^1, pH 7.0).

### Affinity purification of Vhu-Fdh-Hdr complexes

Two 500 ml cultures of *M. maripaludis* MM1265 or Δ*fdh1* Δ*fdh2* expressing *fdh2*-His were grown in 1-liter anaerobic Schott bottles (Chemglass) containing McCas-formate medium with a N_2_:CO_2_ (80:20) headspace pressurized to 100 kPa. Upon reaching an OD_600_ of ∼0.6 (17), cells were anaerobically harvested by centrifugation at 4,000 × g for 10 min. The cell pellet was resuspended in 10 ml binding buffer (100 mM NaCl, 12.5 mM MgCl_2_, 25 mM HEPES (pH 7.5), 10 mM imidazole, 0.5 mM dithionite, and 20 μM flavin adenine dinucleotide). Samples were stored at −80°C for up to 4 weeks before use.

Immobilized metal affinity chromatography (IMAC) was carried out as previously described (17) with all steps in the purification performed anoxically in a room temperature in Coy anaerobic chamber (2 to 3% H_2_, balance N_2_ in the atmosphere). Briefly, the *M. maripaludis* frozen cell paste was brought into the anaerobic chamber, thawed on ice, and transferred to a 15-ml conical tube. The cell suspension was sonicated using a QSonica model XL-2000 sonicator at a setting of 12 for ∼3 seconds with a 10-second cool-down interval on ice for a total of 10 pulses. Upon removal of cellular debris by centrifugation at 16,000 × g for 10 min, 1 ml of nickel-chelating resin slurry (G-Biosciences) was added to the cleared lysate. The resin-lysate suspension was gently mixed by inverting the tube every 10 min for a total of 1 hour. The sample was poured into a disposable gravity flow column (Thermo Scientific) until the resin bed settled. Once the flow through was removed, the column was washed three times with 5 ml wash buffer (same components as binding buffer except with 30 mM imidazole). Finally, proteins were eluted with 2.5 ml elution buffer (same components as binding buffer except with 300 mM imidazole). The eluate was concentrated using Pierce^TM^ Protein Concentrators (10k-MWCO) (Thermo Scientific) by centrifugation for 20 min at 10,000 × g.

The concentrated affinity-purified protein sample underwent further purification by fast performance liquid chromatography (FPLC) using an AKTA purifier UPC10 system (GE Life Sciences) and a Hiprep 16-60 Sephacryl S300HR size exclusion chromatography column at a flow rate of 1 ml min^-1^ (running buffer was the same as the binding buffer but without the imidazole). Specific 1 ml fractions were collected, pooled, and concentrated based on the previously reported retention time of *M. maripaludis* Fdh-Hdr complex for the Hiprep column (21).

The purified Fdh-Hdr complex proteins were analyzed by sodium dodecyl sulfate-polyacrylamide gel electrophoresis (SDS-PAGE) followed by staining with Coomassie blue. The protein samples were also submitted to mass spectrometry analysis and used in *in vitro* biochemical assays to determine the specific enzyme activity with formate as the electron donor.

### Specific activity of Vhu-Fdh-Hdr complexes

The flavin-based electron bifurcation assay for the purified Fdh-Hdr complex was carried out as previously described (17) with the following modifications. Briefly, the assay was carried out in three different conditions based on the presence of potential electron acceptor(s): metronidazole only, F_420_ only, or both. In a Coy anaerobic chamber containing 2-3% H_2_ with balance N_2_, the following reagents were premixed inside a cuvette with a total volume of 487 μl: 1.6 M potassium phosphate (pH7), 700 μM CoB-S-S-CoM, 1 μg of purified Fdh1-Hdr or Fdh2-Hdr complex, and a chosen electron acceptor of either 180 μM metronidazole, or 10 μM F_420_, or both. The cuvette was sealed with a butyl rubber stopper and capped with a screw cap before the headspace was flushed with N_2_ gas for 5 minutes and pressurized to 140 kPa (17).

Using a Cary 3500 UV-visible (UV-Vis) spectrophotometer (Agilent), the cuvette was preheated at 37°C and allowed to equilibrate for 5 min before the reaction was started. The absorbance reading measurement was set at 320 nm if only metronidazole was present, 420 nm if only F_420_ was present, or both 320 nm and 420 nm when both electron acceptors were present. To establish a baseline reading, absorbance was recorded for 2 to 5 minutes without any addition. Then, 20 mM sodium formate was added using a Hamilton syringe to start the reaction. The reaction was monitored for 15 minutes.

The first ∼10% of the reaction curve was used to calculate specific activity of metronidazole or F_420_ reduction (17). The rate of absorbance change (Abs min^−1^) was divided by the extinction coefficient of metronidazole at 320 nm (ε = 9,300 M^−1^ cm^−1^) or of F_420_ at 420 nm (ε = 41400 M^−1^ cm^−1^) (37, 38). The resulting value was multiplied by 0.0005 liters and divided by the amount of protein in the sample (0.001 milligrams) to obtain the specific activity in micromoles per minute per milligram.

### Mass spectrometry analysis

Mass spectrometry was performed with the help of the University of Minnesota Center for Mass Spectrometry and Proteomics. A 5 µg aliquot (10ul) of each sample was made and 50 µg denaturing buffer was added to each sample (7 M urea, 2 M thiourea, 0.4 Tris pH 8, 20% acetonitrile, 10 mM TCEP, and 40 mM chloroacetamide). Samples were incubated at 37°C for 0.5 hrs. 190 µl of LC-MS grade water was added to dilute the urea. Trypsin was reconstituted in 10 mM CaCl_2_ (sequencing grade, Promega, Middleton, WI) and added at a 1:40 enzyme to total protein ratio. Samples were incubated for 16 hrs at 37°C. After incubation, samples were acidified to 0.3% formic acid with 10% formic acid and cleaned up with an MCX STAGE tip (39) using Empore SDB-RPS extraction disks material from 3M. The eluted samples were Speedvaced to dryness.

Analytical separation and detection were performed on an UltiMate 3000 RSLCnano UHPLC system (Thermo Scientific, Waltham, MA) interfaced to an Orbitrap Fusion Tribrid mass spectrometer (Thermo Fisher Scientific, San Jose, CA). All dried peptide samples were reconstituted using a load solvent mixture of 98:2, H_2_O:acetonitrile (ACN) with 0.1% formic acid (FA). 200 nanograms of peptide mixture in 2 µL were injected on the analytical platform equipped with a 10 µL injection loop. Chromatographic separation was performed using a self-packed C18 column (Dr. Maisch GmbH ReproSil-PUR 1.9 µm 120 Å C18aq, 100 µm ID x 45 cm length) maintained at 55 °C for the duration of the experiment. The mobile phases were (A) 0.1% FA in H_2_O and (B) 0.1% FA in ACN solutions. Chromatographic separation was performed using a linear gradient as follows: 5% B from 0-2.5 min, 21% B at 40 min, 35% B at 60 min, and 90% B from 62-69 min followed by a return to starting conditions. The flow rate was operated at 400 nl min^-1^ for 0-2 min, 315 nl min^-1^for 2.5-60 min, and 400 nl min^-1^for 62-69 min. A Nanospray Flex ion source (Thermo Fisher Scientific) was used with a source voltage of 2.1 kV and ion transfer tube temperature of 250 °C.

Discovery LC-MS/MS analyses was performed using full-scan detection followed by data dependent MS^2^ acquisition (DDA). Full-scan detection was performed using Orbitrap detection at a resolution of 120,000, normalized automatic gain control (AGC) targeted setting of 4×10^5^ with a 50 ms maximum ion injection time. Scan ranges of 380 *m/z* – 1580 *m/z* were used for full-scan detection. MS^2^ spectra were collected using a DDA design with a 3 sec cycle time in centroid mode. Fragment spectra were acquired with quadrupole isolation width of 1.6 *m/z*, ion trap detection, and an AGC setting of 1×10^4^ with a 35 ms maximum injection time. The analysis of peptide fragments utilized collision induced dissociation (CID) activation at a constant collision energy of 35%, 0.25 activation Q, and 10 ms activation time.

Peptide tandem MS results were searched and compiled into a protein report using Proteome Discoverer, version 2.5 (Thermo Fisher Scientific, San Jose, CA). Peptide identities were inferred through database searching using the Sequest algorithm against the *Methanococcus maripaludis* (strain S2 / LL) protein database (Proteome ID: UP000000590) downloaded from the online UniProtKB repository (downloaded: 2022-11-17; total protein sequences: 1,722) concatenated to a list of common laboratory protein contaminants (40). The protein database search matched trypsin-specific peptides, maximum 2 missed cleavage sites, with a precursor ion tolerance of 15 PPM and fragment ion mass tolerance of 0.6 Da. Variable peptide modifications included oxidation of methionine, pyroglutamic acid conversion from glutamine, deamidation of asparagine, acetyl and/or met-loss of the protein N-terminus. Carbamidomethylation of cysteine was specified as a fixed peptide modification. Peptide and protein identification were restricted to a 1% false discovery rate using the Percolator algorithm (41).

### Identification of Δ*fdh1* Δ*fdh2* + *fdh2*\* suppressor mutations

The suppressor mutant Δ*fdh1* Δ*fdh2* + *fdh2*\* grown in McCas-formate was incubated until it reached stationary phase. Then, 2 ml of the culture was harvested and centrifuged at 11,000 × g for 10 minutes where the cell pellet was used for genome extraction with the DNeasy Blood & Tissue Kit (Qiagen) protocols. The extracted DNA was submitted for Illumina 2×151bp whole genome sequencing using the NextSeq 2000 platform through Seqcenter. In parallel, the Δ*fdh1* Δ*fdh2* strain grown in McCas with H_2_ was also harvested, genome extracted, and submitted for genome sequencing to serve as control.

Reads were aligned to the reference genome of *Methanococcus maripaludis* strain S2 (accession BX950229 (42)) using Bowtie2’s (v.2.3.4.1) end-to-end, paired alignments with default parameters (43). Aligned sequences were imported into Geneious prime® (v.2021.0.3) and single nucleotide polymorphism (SNP) analysis was performed using the find variations/SNPs function with default parameters except the minimum variant frequency threshold value was adjusted to from 0.25 to 0.15. SNPs were identified for Δ*fdh1* Δ*fdh2* + *fdh2*\* mutants grown in McCas-formate as well as for a Δ*fdh1* Δ*fdh2* strain grown in McCas with H_2_. Suppressor variants were then compared using the feature comparison tool in Geneious prime® to filter variants present in Δ*fdh1* Δ*fdh2* background unrelated to the ability to grow using formate. Additionally, SNPs identified in *fdhA1, fdhB1, fdhA2, fdhB2, upt,* and *mcrB* were manually excluded since these genes were either not present in the genome analyzed or were part of the recombinant pLW40neo construct.

Initial genome analysis of Δ*fdh1* Δ*fdh2* + *fdh2*\* grown in McCas-formate identified a null mutation in *moeA*. We initially hypothesized that tungsten may replace molybdenum in *fdh2*, so we tested altered concentrations of tungsten to stimulate growth. The final concentration of tungsten (Na_2_WO_4_.2H_2_0) in these experiments was increased to 2 × 10^-4^ %, 4 × 10^-4^ %, 6 × 10^-4^ %, 8 × 10^-4^ %, and 1 × 10^-3^ % (wt/v). The Δ*fdh1* Δ*fdh2* + *fdh2* strain was sub-inoculated into these modified McCas-formate media and unmodified McCas-formate as control with duplicates. In parallel, the WT strain was also grown. We saw no differences in growth under any condition tested, so altered tungsten was not considered further. After growth, the Δ*fdh1* Δ*fdh2* + *fdh2*\* strains were harvested, DNA was extracted, and submitted for genome sequencing as described above. The same DNA reads analysis was carried out for these samples.

### RNAseq analysis

For RNA extraction, 1 ml of WT and Δ*moeA* stationary culture grown in McCas-formate was used with the Direct-zol RNA Miniprep Kits protocols. Purified RNA was submitted to Sequencing Center for analysis. Quality control and adapter trimming was performed with bcl-convert. Read mapping was performed with HISAT2 (44). Read quantification was performed using Subread’s featureCounts functionality (45). Read counts were loaded into R (46) and were normalized using edgeR’s (47) Trimmed Mean of M values (TMM) algorithm. Subsequent values were then converted to counts per million (cpm). Differential expression analysis was performed using edgeR’s Quasi-Linear F-Test (qlfTest) functionality against treatment groups. The results of the qlfTest for all genes in addition to the normalized counts per million were generated. Based on this, is a subset of the data with |logFC| > 1 and p < .05 were selected as the genes with the highest RNA log fold-changes.

## Results

### Fdh2 can substitute for Fdh1 in supporting formate-dependent growth in *M. maripaludis*

*M. maripaludis* possesses two formate dehydrogenases. Prior studies found that Fdh1 is essential for growth on formate (19) while Fdh2 may be important under growth conditions when formate concentrations limit growth (31). Interestingly an *M. maripaludis* Δ*fdh1* mutant strain eventually grows on formate after extended incubation (23), suggesting that Fdh2 is sufficient for growth on formate. We hypothesize that the Δ*fdh1* mutant strain acquires a suppressor mutation that allows of Fdh2 to function under these conditions.

To test this hypothesis, we complemented a Δ*fdh1* Δ*fdh2* mutant with a plasmid harboring *fdh2*; this construct contained a C-terminal histidine tag on FdhA2 to facilitate downstream purification. Like Δ*fdh1*, this strain required extended incubation before growth was observed in formate containing medium. Growth of wild-type was observed ∼5 hours after inoculation, whereas the Δ*fdh1* Δ*fdh2* strain expressing the *fdh2-His* construct only began to grow ∼35 hours after inoculation (Figure 2). When subcultured into fresh medium, this strain had a reduced lag period, suggesting that a suppressor mutation was responsible for the observed growth. We call this strain Δ*fdh1* Δ*fdh2* + *fdh2*\*.

**Figure 2:**
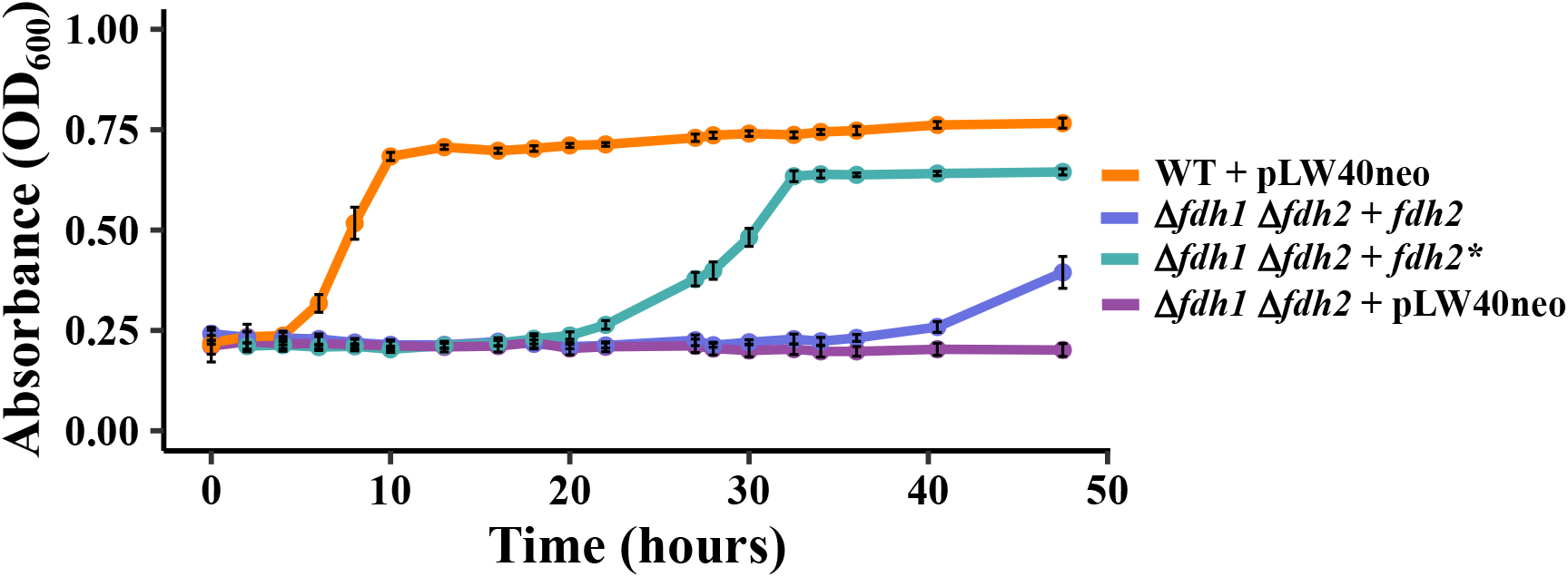
Cells expressing *fdh2* are able to grow with formate as sole electron donor after an extended lag phase. Growth of *M. maripaludis* wild-type, Δ*fdh1* Δ*fdh2* + pLW40neo, Δ*fdh1* Δ*fdh2* + *fdh2*, Δ*fdh1* Δ*fdh2* + *fdh2*\* in McCas-formate medium. Data are averages and standard deviations from quadruplicate cultures. There was a significant lag phase for the *fdh2* expressing cells but upon sub-inoculation, the *fdh2*\*-bearing suppressor mutations have a reduced lag phase.

### Both Fdh1 and Fdh2 are associated with heterodisulfide reductase

Because the Δ*fdh1* Δ*fdh2* + *fdh2*\* strain grows on formate, we sought to determine whether Fdh2 performs similar functions as Fdh1. In wild-type, Fdh1 associates with Hdr and facilitates electron transfer from formate (21). Leveraging the His-tag on FdhA2, we purified the protein by immobilized metal affinity chromatography. A Coomassie-stained SDS-PAGE gel (Figure 3) suggested that, like Fdh1, Fdh2 associates with Hdr (20, 21). The number of bands and their size is consistent with a protein complex composed of Fdh2, Hdr, and the F_420_-non-reducing hydrogenase (Vhu).

**Figure 3:**
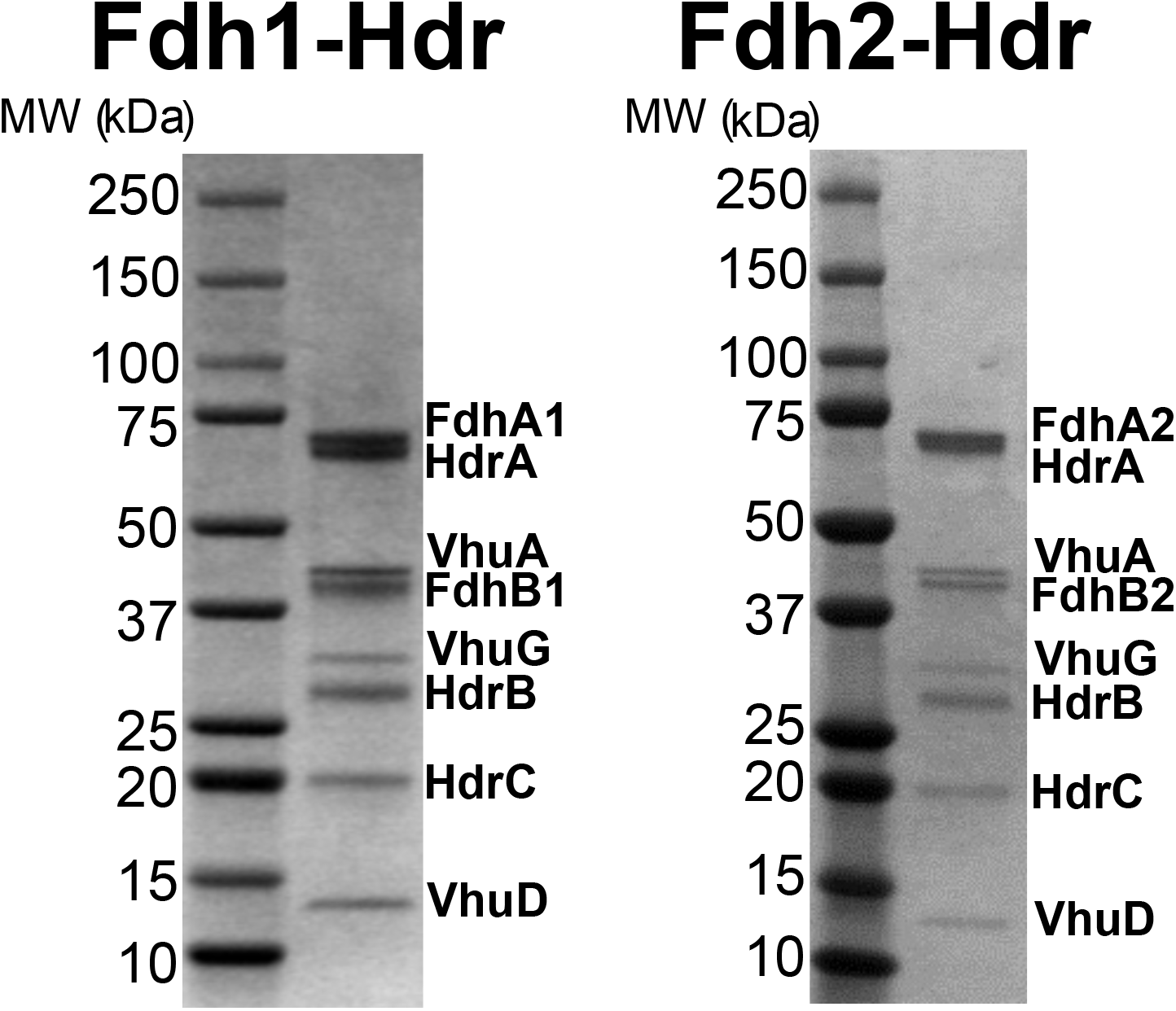
Coomassie-stained SDS-PAGE gel of purified Vhu-Fdh-Hdr complex of Δ*fdh1* Δ*fdh2* expressing *fdh1-His* (A) or *fdh2-His* (B).

To confirm the composition of the protein complex, we submitted the protein for mass spectrometry analysis. FdhA2 and FdhB2 were both present in samples prepared from the Δ*fdh1* Δ*fdh2* + *fdh2*\* strain, indicating successful heterologous expression of *fdh2* (Table 1). Additionally, HdrA, HdrB2, HdrC2, VhuA and VhuD were present, verifying that Fdh2 interacts with the native Hdr-Vhu protein complex, consistent with purification of heterogenous Vhu-Fdh-Hdr protein complexes produced by *M. maripaludis* (20).

**Table 1:** Proteins identified by mass spectrometry analysis of affinity-purified protein samples from *M. maripaludis* Δ*fdh1* Δ*fdh2* expressing *fdh2\**

| Description | Gene Symbol | MW in kDa | Coverage | Total number of PSMs |
| --- | --- | --- | --- | --- |
| H<sub>2</sub>/formate:CoB-CoM heterodisulfide,ferredoxin reductase subunit A | *hdrAu* | 71.2 | 57% | 848 |
| Formate dehydrogenase alpha subunit | *fdhA2* | 74.7 | 66% | 778 |
| F<sub>420</sub>-non-reducing hydrogenase subunit alpha | *vhuA* | 45.6 | 92% | 613 |
| H<sub>2</sub>/formate:CoB-CoM heterodisulfide,ferredoxin reductase subunit C2 | *hdrC2* | 20.4 | 75% | 464 |
| H<sub>2</sub>/formate:CoB-CoM heterodisulfide,ferredoxin reductase subunit B2 | *hdrB2* | 32 | 57% | 371 |
| Formate dehydrogenase beta subunit | *fdhB2* | 42.4 | 59% | 342 |
| Chaperonin GroEL (Thermosome, HSP60 family) | *hsp60* | 58.2 | 69% | 294 |
| Formylmethanofuran dehydrogenase, subunit A | *fmdA* | 64.6 | 79% | 238 |
| Coenzyme F<sub>420</sub>-reducing hydrogenase subunit alpha | *fruA* | 46.2 | 75% | 235 |
| H<sub>2</sub>/formate:CoB-CoM heterodisulfide,ferredoxin reductase subunit B1 | *hdrB1* | 33.5 | 47% | 220 |
| F<sub>420</sub>-non-reducing hydrogenase vhu iron-sulfur subunit D | *vhuD* | 14.7 | 49% | 217 |
| Acetolactate synthase | *ilvB* | 63.8 | 68% | 164 |
| Tungsten containing formylmethanofuran dehydrogenase, subunit F | *fwdF* | 38.2 | 28% | 144 |
| Phosphoenolpyruvate synthase | *ppsA* | 83.3 | 64% | 131 |

### The Vhu-Fdh2-Hdr complex exhibits similar F_420_ and metronidazole reduction activities as the Fdh1-containing complex

Using purified Vhu-Fdh2-Hdr, we sought to test the ability of this complex to oxidize formate concomitant with either F_420_ reduction or the flavin-based electron bifurcating activity of Hdr (48). To determine how Fdh2 activity compares to Fdh1 activity, we also performed all assays using Vhu-Fdh1-Hdr protein complexes. The tail of F_420_ can vary widely between different species of methanogenic archaea (49); therefore, F_420_ was purified from *M. maripaludis* following established protocols (50). For the activity measurements, F_420_ reduction was monitored based on a decrease in absorbance at 420 nm, and metronidazole reduction was monitored based on a decrease in absorbance at 320 nm. We used metronidazole as an alternate electron acceptor to ferredoxin (Fd) to avoid overlapping spectral signals between F_420_ and Fd (abs at 390 nm vs. 320 nm for metronidazole). All assays containing metronidazole also contained CoM-S-S-CoB to facilitate the electron bifurcating reaction.

Similar to the Vhu-Fdh1-Hdr complex, the Vhu-Fdh2-Hdr complex exhibited robust F_420_ reduction activity (Figure 4). Next, we tested the flavin-based electron bifurcation activity of both Vhu-Fdh-Hdr complexes with formate as the electron donor. Upon formate addition, metronidazole reduction occurred for the Vhu-Fdh2-Hdr complex suggesting that Fdh2, like Fdh1 (20), can facilitate formate oxidation coupled to the electron bifurcating activity of Hdr.

**Figure 4:**
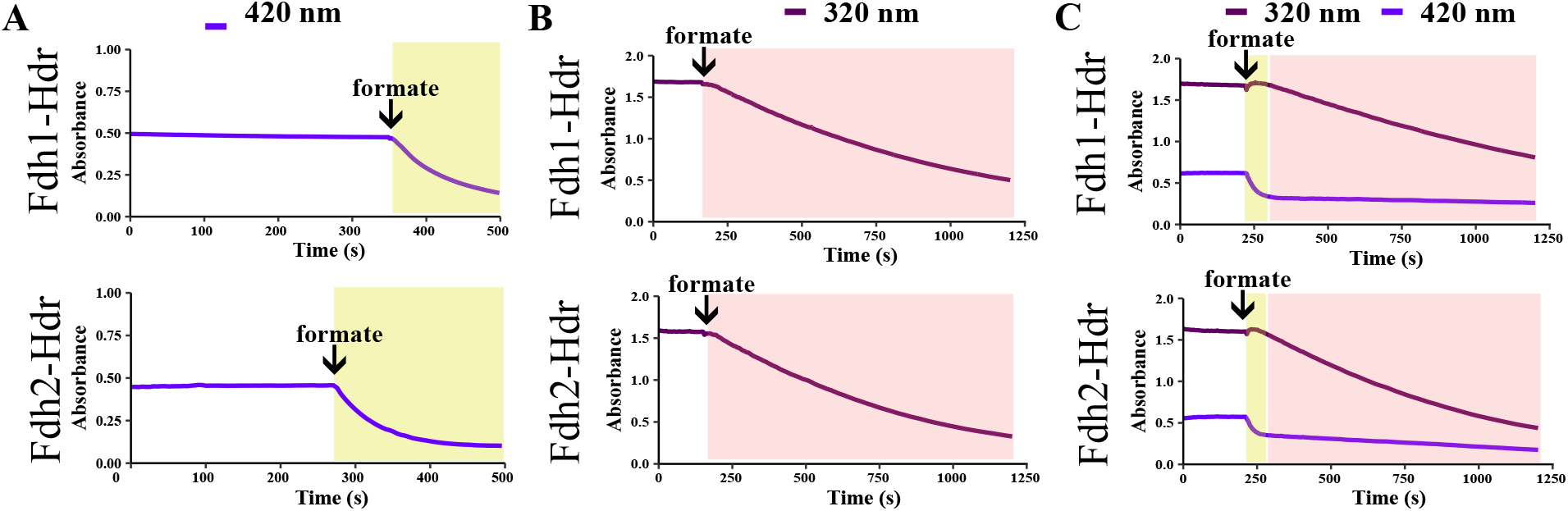
Fdh1 and Fdh2 exhibit both F_420_-reduction activity and mediate electron transfer for electron bifurcation by Hdr. (A) F_420_ reduction by purified Vhu-Fdh-Hdr complexes. (B) metronidazole reduction by purified Vhu-Fdh-Hdr complexes. (C) F_420_ and metronidazole reductions catalyzed by Vhu-Fdh-Hdr complexes. F_420_ reduction was tracked by absorbance change at 420 nm and metronidazole reduction was tracked by absorbance change at 320 nm. F_420_ reduction and metronidazole reduction reactions are highlighted with yellow and pink backgrounds, respectively.

Because both the Vhu-Fdh1-Hdr and Vhu-Fdh2-Hdr complexes are able to independently reduce F_420_ or metronidazole coupled to formate oxidation, we sought to determine whether one electron flow path is preferred over another. F_420_ (monitored at 420 nm), metronidazole (monitored at 320 nm) and CoM-S-S-CoB were all present in the reaction mixture, and the reaction was initiated by the addition of formate. Upon formate addition there was an immediate reduction in absorbance at 420 nm. Absorbance at 320 nm only changed after the F_420_ reducing reaction proceeded to completion (Figure 4). This pattern was observed for both Vhu-Fdh1-Hdr and Vhu-Fdh2-Hdr complexes, indicating that the formate dependent reactions reduce F_420_ before initiating the electron bifurcation reaction, regardless of which Fdh is associated with Hdr.

Using the reaction data, we determined the specific activity for F_420_ and metronidazole reduction for both the Fdh1 and Fdh2 containing protein complexes. For F_420_ reduction, the Fdh1-containing complex exhibited a specific activity of 10.11 μmol mg^-1^ min^-1^ while Fdh2-containing complex exhibited an average specific activity of 10.21 μmol mg^-1^ min^-1^ (Figure 5). For metronidazole reduction, the Fdh1-containing complex has a specific activity of 9.71 μmol mg^-1^ min^-1^ and the Fdh2-containing complex has a specific activity of 9.88 μmol mg^-1^ min^-^. When both F_420_ and metronidazole were provided, the Fdh1-containing complex exhibited a specific activity of F_420_ reduction of 10.59 μmol mg^-1^ min^-1^ and a specific activity of metronidazole reduction of 10.33 μmol mg^-1^ min^-1^, and the Fdh2-containing complex exhibited a specific activity of F_420_ reduction of 8.74 μmol mg^-1^ min^-1^ and a specific activity of metronidazole reduction of 10.25 μmol mg^-1^ min^-1^ (Figure 5). These results indicate that both Fdh1 and Fdh2 have similar specific activities and likely perform similar functions in the cell. Nonetheless, it is still unclear why Fdh1, but not Fdh2, is critical for growth on formate. Our growth data of the Δ*fdh1* Δ*fdh2* + *fdh2*\* strains suggested that a suppressor mutation could be responsible for altering the expression or activity of Fdh2.

**Figure 5:**
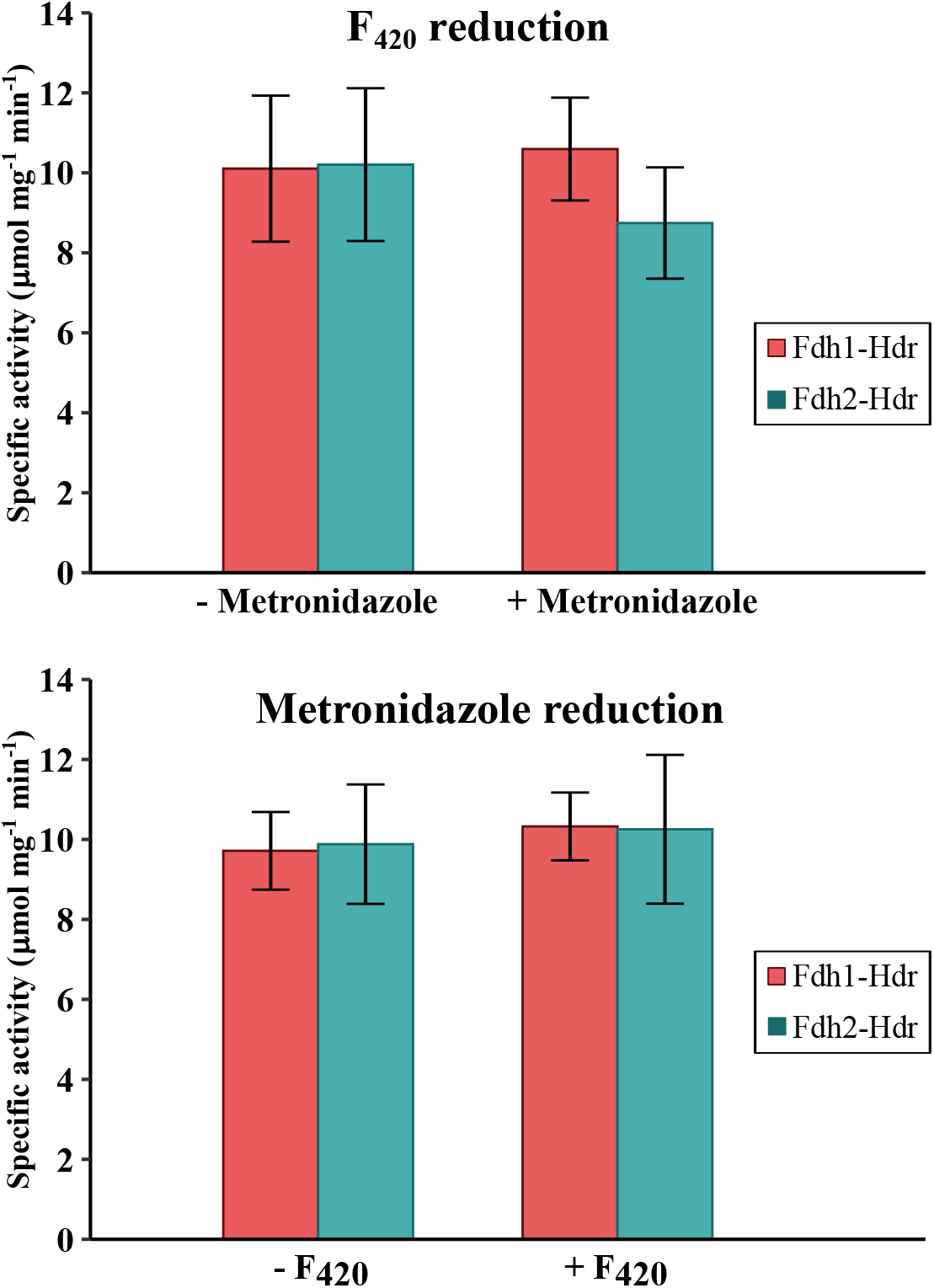
Both Vhu-Fdh-Hdr protein complexes exhibit a similar specific activity for the reduction of F_420_ and metronidazole. Data are from triplicate measurements and error bars represent one standard deviation around the mean. There were no significant differences in activity (Student’s *t*-test, p > 0.05).

### Mutations in *moeA* are common in Δ*fdh1* Δ*fdh2* + *fdh2*\* strains

Because the growth pattern of cells relying on *fdh2* expression for growth suggested a suppressor mutation occurred, we sought to characterize this mutation. We initially hypothesized that mutations in the *fdh2* gene itself allowed for growth. To test this, the expression plasmid was extracted from suppressor mutant strains and sequenced. There were no mutations in *fdh2* or in other regions of plasmid, suggesting that a mutation occurred on the genome. We isolated 11 independently generated Δ*fdh1* Δ*fdh2* + *fdh2*\* suppressor strains and sequenced genomic DNA to identify mutations. Mutations were identified in 7 out of 11 of these strains and the majority of suppressor mutants contained mutations in a molybdopterin transferase (*moeA*, locus tag: MMP_RS08320). Across multiple strains, we identified two mutant alleles for this gene; a nonsense mutation in the codon for Ser^132^ and an adenine base deletion in the codon encoding Lys^150^ that led to a frameshift mutation. Due to the high number of suppressor strains containing mutated *moeA*, we focused our efforts on that gene. Other mutations were identified (Figure 6), but they never occurred in more than one independent strain so were not considered further.

**Figure 6:**
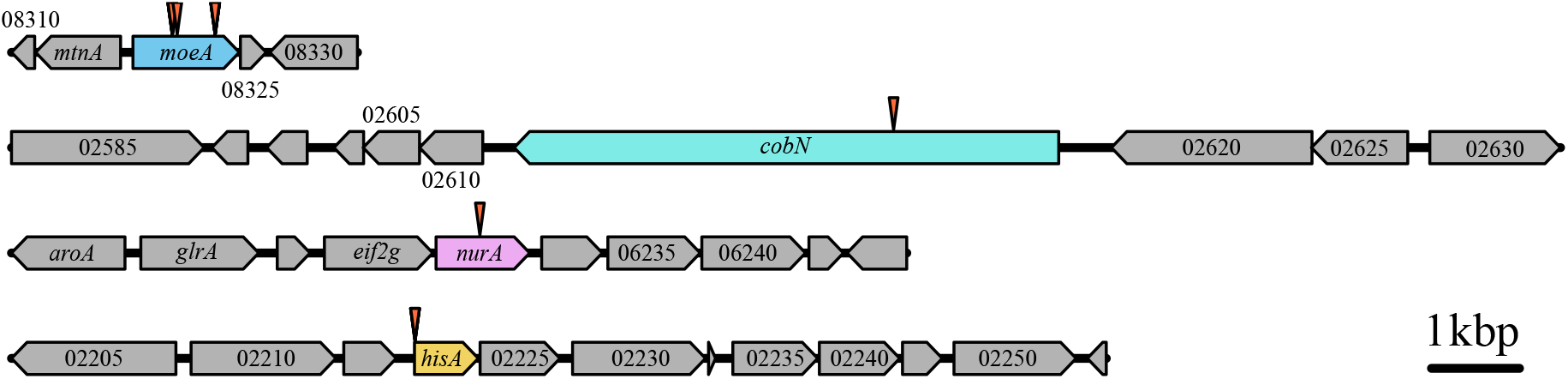
Genomic location of mutations identified in Δ*fdh1* Δ*fdh2* + *fdh2*\* strains. A total of four distinct genes sustained mutations across the 11 sequenced isolates. Among these, multiple alleles were found for *moeA*, while single mutations were found in the *cobN*, *nurA*, and *hisA*. Other unannotated genes are referenced using the locus tag format of MMP_RS0XXXX.

### Deletion of *moeA* alters *fdh* expression and is sufficient for *fdh2*-dependent growth

To validate the suppressor analysis, we generated a mutant lacking *moeA* in the *Δfdh* background (Δ*fdh1* Δ*fdh2* Δ*moeA*) and reintroduced recombinant plasmid containing either *fdh1* or *fdh2*. The Δ*fdh1* Δ*fdh2* Δ*moeA* + *fdh1* strain was unable to grow in formate containing medium while growth was rescued in the Δ*fdh1* Δ*fdh2* Δ*moeA* + *fdh2* background to a level observed in suppressor strains (Figure 2 and Figure 7). The lack of growth in the Δ*fdh1* Δ*fdh2* Δ*moeA* + *fdh1* strain suggests that *moeA* is responsible for synthesis of the molybdopterin cofactor of Fdh1, and lack of this gene effectively generates an *fdh1* null phenotype. The *fdh1* expression plasmid encoded a functional copy of *fdh1* because growth was rescued in genetic backgrounds that contained *moeA* (Δ*fdh1* Δ*fdh2* + *fdh1*) (Figure 7).

**Figure 7:**
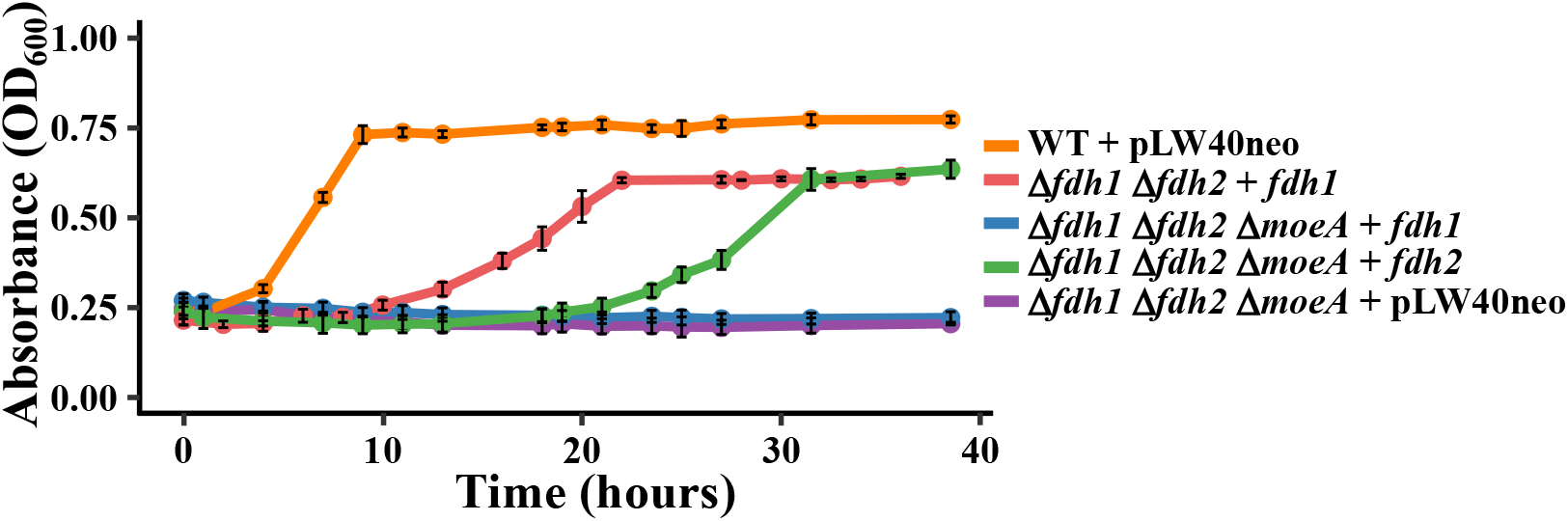
Deletion of *moeA* is sufficient for growth of *fdh2-His* expressing cells when formate is the sole electron donor.

Loss of Fdh1 function in Δ*moeA* mutant strains, coupled to the activity of Fdh2, suggested that regulation of *fdh2* is altered in this background. We generated a *moeA* (MMP_RS08320) deletion strain (Δ*moeA*) and submitted it for RNAseq in order to determine if this mutant background had altered expression of genes encoding Fdh homologs. Indeed, when the Δ*moeA* strain was compared against the WT strain, transcripts of *fdh2* were significantly elevated (Table 2). We did not note a difference in *fdh1* mRNA abundance. We also found an increase in mRNA abundance for genes encoding EhaD, FmdB, FlaB1, and another MoeA homolog (MMP_RS02715), suggesting that a different *moeA* gene may be responsible for synthesis of the molybdopterin cofactor of Fdh2.

**Table 2:** Genes with the highest RNA log_2_ fold-changes (log_2_FC) in Δ*moeA* compared to WT.

| Locus Tag | Gene Name | Description | Log<sub>2</sub>FC |
| --- | --- | --- | --- |
| MMP_RS00800 | *fdhB2* | formate dehydrogenase beta subunit | 7.016173 |
| MMP_RS07465 | *ehaD* | energy-conserving hydrogenase delta subunit | 5.111884 |
| MMP_RS00795 | *fdhA2* | formate dehydrogenase alpha subunit | 4.954532 |
| MMP_RS08485 | *NA* | molybdate ABC transporter, substrate binding subunit | 4.331484 |
| MMP_RS02705 | *fmdB* | molybdenum containing formylmethanofuran dehydrogenase, subunit B | 4.329776 |
| MMP_RS08580 | *flaB1* | flagellin B1 precursor | 4.180443 |
| MMP_RS03010 | *NA* | conserved hypothetical protein | 3.973806 |
| MMP_RS07455 | *NA* | conserved hypothetical protein | 3.929334 |
| MMP_RS04300 | *NA* | hypothetical protein | 3.763632 |
| MMP_RS02715 | *moeA* | molybdopterin biosynthesis protein | 3.751437 |

## Discussion

Methanogens that grow with formate as an electron donor possess multiple isoforms of Fdh on their genome. In *M. maripaludis*, Fdh1 is the primary isoform, donates electrons to both F_420_ and Hdr, and is required for growth in batch culture (20, 23, 31). *fdh1* is expressed when H_2_ concentrations limit growth or when formate is provided as a sole electron donor, and *fdh2* is expressed under conditions where formate concentrations limit growth (31, 51). Thus, multiple isoforms of Fdh are thought to allow the methanogens to adapt to different environments, but it is unclear whether the multiple isoforms are functionally redundant with regards to the reactions they catalyze. To determine the roles of multiple Fdhs in methanogenesis, we sought to determine the activity of Fdh1 and Fdh2 in *M. maripaludis* and found that both enzymes catalyze the same reactions with similar specific activities, but they may be regulated such that they are only expressed under specific growth conditions.

Fdh is required for two distinct reactions in different steps of methanogenesis: F_420_ reduction (52–54) and electron transfer to the Hdr complex (20). However, what is not clear is whether these activities are coordinated. To provide all necessary reducing equivalents for the reduction of one CO_2_ to CH_4_, four electrons are needed to reduce two molecules of F_420_, and four electrons are required for the electron bifurcating reaction that reduces ferredoxin and CoM-S-S-CoB (20, 55). Interestingly, we found that the Hdr-bound form of Fdh exhibits remarkable versatility by performing all of these reactions, with a preference for F_420_ reduction as the initial step *in vitro* (Figure 4). In our purifications, Fdh was histidine tagged and targeted for purification, and based on densitometry analysis, all recovered protein was Hdr associated. This suggests that unbound Fdh is not abundant in whole cells. Interestingly, each isoform of Fdh appears to have dedicated machinery for synthesis of the molybdopterin cofactor. This is apparent from the increased expression of a gene encoding a second isoform of MoeA in the Δ*moeA* mutant background and the fact that Fdh1 is non-functional when the putative MoeA isoform responsible for its molybdopterin cofactor synthesis is absent. We hypothesize that the competing, but identical, reactions catalyzed by Fdh1 and Fdh2 are regulated partly in response to synthesis of these cofactors.

The observation that F_420_ reduction proceeded to completion before initiating of the flavin-based electron bifurcation reaction is likely due to an excess of formate provided in the enzyme assay. In most methanogens, the cellular ratio of oxidized to reduced F_420_ varies with the concentration of electron donors in the environment, where this ratio decreases when excess reductant is present (56). Therefore, we hypothesize that the competing reactions, where both F_420_ and metronidazole are present, would favor F_420_ reduction due to excess formate in the reaction mixture. Alternatively, a preference for F_420_ reduction may be a necessity given the anabolic needs of the cell requiring H_2_ (24, 57). When cells are grown with formate, F_420_H_2_ is an electron donor for H_2_ production by the F_420_-reducing hydrogenase or by the combined activities of the F_420_-dependent methylenetetrahydromethanopterin dehydrogenase and the H_2_-dependent methylenetetrahydromethanopterin dehydrogenase (34). Prioritizing F_420_ reduction to maintain high a concentration of F_420_H_2_ ensures cells can meet their biosynthetic demands.

While H_2_ metabolism is widespread among methanogens, a growing body of literature highlights the importance of formate as an electron donor, and an expanded understanding of the biochemistry of formate oxidation is necessary to optimize methanogenic metabolism. Methanogens from the *Methanomicrobiales* may be formate specialists because the essential reduction of CoM-S-S-CoB appears to require formate, but not H_2_, in these organisms (17, 18). Furthermore, *in situ* growth conditions, especially when methanogens grow in syntrophic association with partner bacteria, may favor formate-dependent metabolism in many environments (16, 58, 59). Our results suggest that methanogens that encode multiple isoforms of Fdh possess redundancy to optimize activity under a variety of growth conditions.

## Acknowledgements

We thank William Whitman for providing cell pellets used for the purification of F_420_. This work was funded by the U.S. Department of Energy, Office of Science, Basic Energy Sciences under grant DE-SC0019148.

## Notes

### Competing Interest Statement

The authors have declared no competing interest.

